# How Do Telomeres Block Checkpoint Activation?

**DOI:** 10.1101/251389

**Authors:** Julien Audry, Jinyu Wang, Jessica R. Eisenstatt, Steven L. Sanders, Kathleen L. Berkner, Kurt W. Runge

## Abstract

Genome instability is a potentially lethal event for a eukaryotic cell, and a mutational force for genetic diseases such as cancer. DNA double-strand breaks (DSBs) can drive genome instability and are sensed by the DNA damage checkpoint, a defined set of evolutionarily-conserved proteins that bind the DSB to signal a pause or arrest of the cell cycle^1^ and recruit proteins to repair the DNA lesion^2,3^. Telomeres, the physical ends of linear eukaryotic chromosomes, are specialized DSBs that suppress DNA damage checkpoint activation by an unknown mechanism(s), even though telomeres are bound by many of the DNA damage checkpoint proteins that signal cell cycle arrest^4^. Carneiro et al. (*Nature* 467: 228-232) addressed this question using *Schizosaccharomyces pombe* cells that lack Taz1, the protein that binds to double-stranded telomere repeats^5^. Telomeres in *taz1*∆ cells have single-stranded DNA regions that are bound by checkpoint and DNA repair proteins, but cells do not arrest^5,6^. Carneiro et al. found that the ortholog of the human DNA damage checkpoint protein 53BP1 (Crb2) found at DSBs was not recruited to telomeres^5^. Crb2 can bind to dimethylated lysine 20 of histone H4 (H4K20me2) in nucleosomes^7^. Carneiro et al. presented data that H4K20me2 was depleted near telomeres in wild type and *taz1*∆ cells, suggesting a mechanism for checkpoint suppression^5^. Our efforts to pursue this exciting result led to the discovery that H4K20me2 is not depleted near telomeres, indicating that checkpoint suppression occurs by some other mechanism(s).

H4K20me2 constitutes ∼25% of total H4 in *S. pombe*^8^, implicating a telomere-associated demethylase to deplete H4K20me2 at telomeres in *taz1*∆ cells to prevent checkpoint-mediated arrest. However, both a genome-wide screen of gene deletion mutants (D. Durocher, pers. comm.) and our screen of demethylase mutants failed to identify a mutant that caused *taz1*∆ cells to arrest, to grow poorly or to recruit more Crb2. We therefore re-evaluated H4K20me2 levels by chromatin immunoprecipitation (ChIP). We first identified commercial antibodies specific for H4K20me2 by western analysis using 11 different samples. Positive controls were extracts from wild type cells, cells where the single *S. pombe* H4K20 methylase gene *set9* is marked and functional (*set9-kan-wt*) and recombinant H4 where the only modification is a chemical mimetic for K20me2^9^. Negative controls included recombinant H4 where the only modifications were mimetics of 0, 1 or 3 methyl groups on lysine 20, and extracts from cells where all three copies of the H4 gene have lysine 20 mutated to arginine (H4K20R). A series of *set9* mutants that methylate H4K20 to contain 0 (*set9*∆), 1 (*set9-F164Y, set9-F178Y*), or 1 and 2 methyl groups (*set9-F195Y*) were also assayed^10^. The specific antibody identified (Fig. 1A) was used in ChIP to monitor H4K20me2 at the telomeric loci assayed in Carneiro et al. and two internal chromosomal loci in wild type and *taz1*∆ cells, and in mutants that lack H4K20 methylation, *set9*∆ and H4K20R. Total H4 levels at these loci were monitored with an antibody that recognizes all H4 forms.

**Figure 1.**
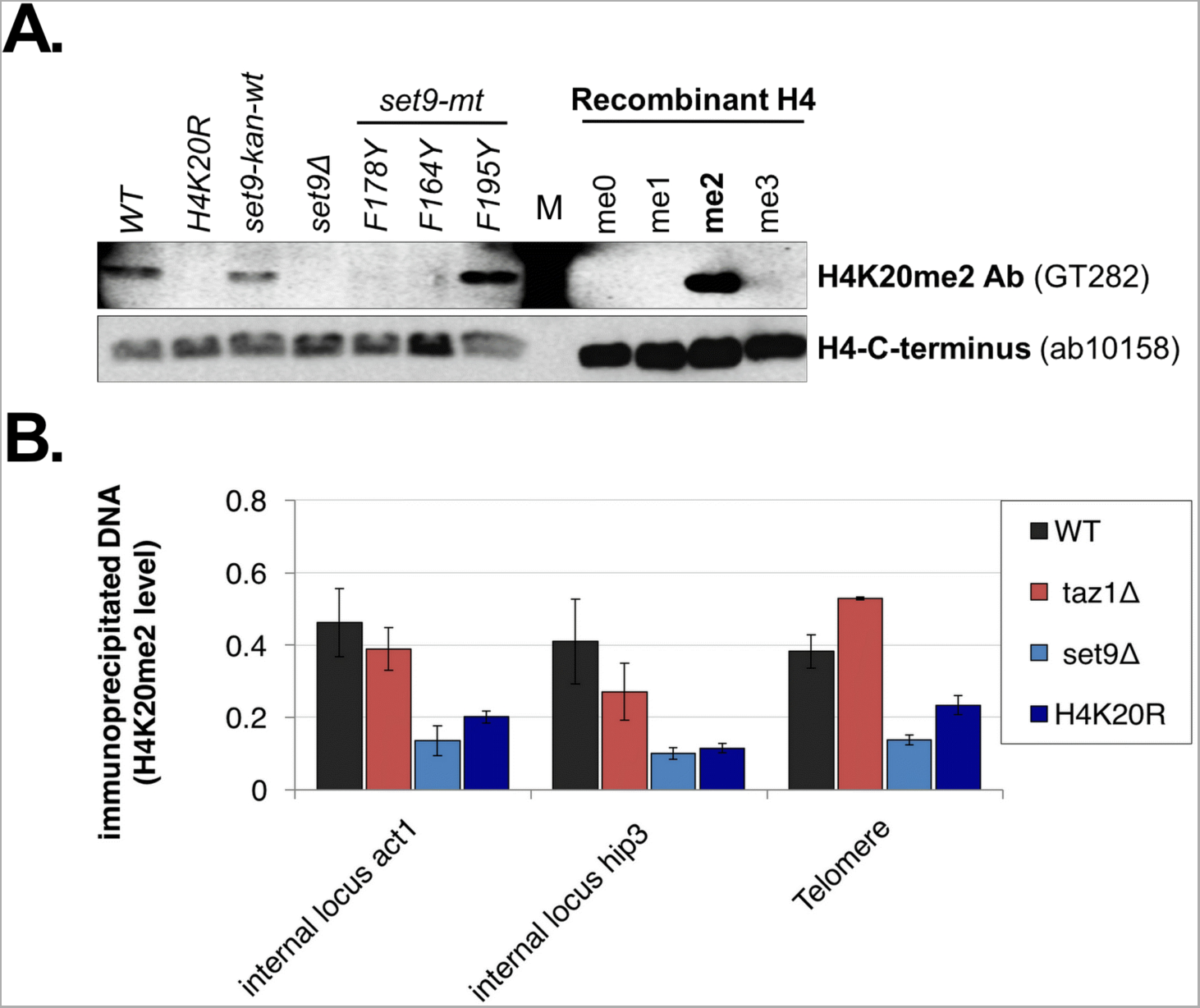
H4K20me2 is not excluded from the telomere repeat-adjacent nucleosomes in wild type or *taz1*∆ cells. An antibody that specifically recognizes H4K20me2 was identified (A) and used in ChIP to measure levels of H4K20me2 in chromatin at two standard internal loci and loci adjacent to the telomere repeats (B). H4K20me2 levels are expressed as the ratio of the H4K20me2 ChIP levels (% of DNA in anti-H4K20me2 IP compared to input chromatin) over H4 ChIP levels (% of DNA in anti-histone H4 IP compared to input chromatin). Wild type and *taz1*∆ cells have the same levels at all loci and are clearly distinguishable from the negative controls (*P* values compared to wild type levels: all *taz1*∆ strains >0.18; all *set9*∆ and H4K20R strains <0.023, individual values are presented in Table S3). Each western blot in panel A used a separate, identically run gel to analyze the samples shown. M stands for molecular weight markers.

We found that H4K20me2 levels are similar at all three loci in wild type and *taz1*∆ cells, and clearly distinguishable from the *set9*∆ and H4K20R negative controls (Fig. 1B). These data argue that while the damaged telomeres in *taz1*∆ cells block checkpoint activation, the mechanism that hinders Crb2 recruitment is unlikely to be the suppression of H4K20me2 in telomeric chromatin. These results and conclusion are consistent with the genetic screen results that did not identify a demethylase required to sever the checkpoint in *taz1*∆ cells, and suggest that searches for combinations of demethylase mutants that activate the checkpoint in *taz1*∆ cells will not be fruitful. Rather, broader approaches to investigate the differences between telomeres and DSBs may be required, including much more extensive characterization of the post-translation modifications of proteins at or near telomeres. While H4K20me2 levels are not reduced at telomeres, it is worth noting that checkpoint activation is the sum of multiple protein interactions and modifications, e.g. phosphorylation of histone H2A and modification of several checkpoint proteins^11,12^. Reducing the efficiency of several of these interactions may be sufficient to impair checkpoint signaling at *taz1*∆ cell telomeres. Results from such studies may provide an understanding of the anti-checkpoint activity of telomeres so that it may be modulated to treat telomere-related diseases such as cellular aging and cancer^13^.

## Supplemental information

accompanies this Comment.

### Author Contributions.

JA, JW, KWR and KLB designed the experiments. JW obtained or made the strains and tested the recombinant H4 histones and then used these reagents to perform the western blots with several commercial antibodies to identify one specific for H4K20me2. JA obtained or created strains for the ChIP experiments, designed the primers and performed these experiments. JRE and SLS designed the experiments to create the H4K20R strains. JRE made and verified the strain and tested it for increased sensitivity to DNA damage (not shown). JA, JW, JRE, KLB and KWR wrote the manuscript.

## Competing Financial Interests

Declared none.

